## Supplementary Information for "HATCHet2: clone- and haplotype-specific copy number inference from bulk tumor sequencing data"

**S4 Haplotype switch correction algorithm 3**

### S1 Benchmarking HATCHet2 against HATCHet [1], Battenberg [2], cloneHD [3], and TITAN [4]

#### S1.1 Overview

Simulated data was generated using the MASCoTE simulator [1] which simulates sequencing reads from *in silico* mixtures of synthetic tumor genomes. We used the 8 multi-sample datasets included in the HATCHet publication [1], which include four datasets with 3 bulk samples each and four with 5 bulk samples each. Each sample is a different mixture of the 2-3 tumor genomes (i.e., clones) simulated for that individual. Half of the datasets include a whole-genome duplication affecting all tumor clones.

#### S1.2 Metrics for evaluating performance on simulated data

We used several different metrics to quantify different aspects of performance in recovering CNAs from simulated data. Let $\mathcal{S}_s$ indicate the set of allele-specific copy-number states that a method infers for a particular genomic segment $s$ across the $n$ tumor clones:

$$\mathcal{S}_s = \{(a_{s,i}, b_{s,i}) | 1 \leq i \leq n\} \tag{S0.1}$$

Let $\overline{\mathcal{S}}_s$ indicate the true copy-number states for region $s$. Let $\ell_s$ indicate the length of genomic segment $s$, and let $L = \sum_s \ell_s$ be the total length of all genomic segments. Then, the precision, recall, and accuracy of allele-specific copy-number states is as follows:

$$\text{precision}(\mathcal{S}_s) = \frac{1}{L} \sum_s \ell_s \frac{|\mathcal{S}_s \cap \overline{\mathcal{S}}_s|}{|\mathcal{S}_s|} \tag{S0.2}$$

$$\text{recall}(\mathcal{S}_s) = \frac{1}{L} \sum_s \ell_s \frac{|\mathcal{S}_s \cap \overline{\mathcal{S}}_s|}{|\overline{\mathcal{S}}_s|} \tag{S0.3}$$

$$\text{accuracy}(\mathcal{S}_s) = \frac{1}{L} \sum_s \ell_s \frac{|\mathcal{S}_s \cap \overline{\mathcal{S}}_s|}{|\mathcal{S}_s \cup \overline{\mathcal{S}}_s|} \tag{S0.4}$$

Following [1], we compute the average allele-specific error per genome position (AASEGP) by considering the clone mixture proportions as a probability distribution over discrete copy-number states and computing the total variation distance between the true and inferred distributions. Specifically, let $u_{s,c,p}$ indicate the proportion of cells in sample $p$ that are assigned allele-specific copy-number state $c$ in segment $s$. Let $\overline{u}_{s,c,p}$ indicate the true value of the corresponding quantity. Let $P$ indicate the number of samples. Then,

$$AASEGP(U) = \frac{1}{LP} \sum_s \sum_{p=1}^{P} \ell_s \max_{c \in \mathcal{S}_s \cup \overline{\mathcal{S}}_s} |u_{s,c,p} - \overline{u}_{s,c,p}| \tag{S0.5}$$

The results reported for Battenberg [2], cloneHD [3], and TITAN [4] on these simulated datasets are the same as those reported in the original HATCHet publication [1].

### S2 Calculation of mean squared error for HATCHet2 and Battenberg

For each method, we computed the mean squared error (MSE) for both BAF and RDR by comparing the expected RDR and BAF of each method's solution (a function of the assigned copy-number states as well as the inferred sample purity and tumor genome length) to the observed BAF and RDR values within the region, taking the difference, and squaring it. Specifically, we evaluated the BAF error at every SNP position in the region and the RDR error at every interval between SNP positions – i.e., given SNPs at positions $i$ and $j$, we computed the RDR in the genomic interval $[i, j)$ in each sample and evaluated the MSE comparing this RDR to the expected RDR.

### S3 Reference-based phasing

#### S3.1 Implementation details

By default, HATCHet2 downloads a 1000 Genomes Project [5] (phase 3, version 5) reference panel that contains haplotypes in hg19 coordinates. To support the phasing of germline SNPs in hg38 coordinates, HATCHet2 also downloads chain files from the UCSC genome browser (http://genome.ucsc.edu) and uses Picard [6] (https://broadinstitute.github.io/picard/) to liftover between the hg19 and hg38 human reference genome versions, both before and after reference-based phasing. HATCHet2 then filters out multi-allelic SNPs and indels using bcftools [7] and phases the resulting bi-allelic germline SNPs with SHAPEIT2 [8].

#### S3.2 Construction of phasing blocks

Given reference-based phasing output, we first iterate over SNPs and apply a test between pairs of adjacent SNPs. Let $x_i$ indicate the number of reads assigned to haplotype "1" at SNP $i$ and $y_i$ indicate the number of reads assigned to haplotype "0" at SNP $i$ (where assignment to haplotypes is given by the reference-based phasing). For each pair $i, j$ of adjacent SNPs, we test whether the binomial proportion $x_i/(x_i+y_i)$ is different from the proportion $x_j/(x_j+y_j)$ at significance level $\alpha$ (default 0.1) using the normal approximation test. Then, we group SNPs to minimize the total number of phasing blocks such that

1 Phasing blocks are contiguous and non-overlapping along the genome.

2 Adjacent SNPs $i, j$ that were significantly different according to the binomial proportion test are not combined.

3 Adjacent SNPs $i, j$ that are more than a distance of $d$ (default 25 kb) apart are not combined in a phasing block.

4 No more than $c$ SNPs (default 10) are combined in a block.

We solve this problem greedily by iterating over SNPs in each bin and forming maximal phasing blocks subject to the above constraints. We merge the counts for all SNPs in each resulting phasing block to form a "meta-SNP" as described in the main text.

### S4 Haplotype switch correction algorithm

While we infer the relative phase for SNPs in each bin when estimating the mhBAF, the relative phase *across* bins may not be consistent, particularly in bins where both haplotypes are approximately in equal abundance (i.e., the "minor" haplotype for the corresponding section of the genome is somewhat ambiguous). In these regions, the minor haplotype may "switch" between adjacent bins, which would result in mhBAF values being reflected about 0.5 in each sample. To ensure that the minor haplotype is consistent across bins on each chromosome arm, we apply the following algorithm to identify and correct haplotype switches. Let $\hat{f}_{s,p}$ indicate the inferred mhBAF value for bin $s \in [1, N]$ and sample $p \in [1, P]$ on a chromosome arm with $N$ bins.

1 Identify samples $p$ for which the average allelic imbalance $\frac{1}{N}\sum_{s=1}^{N} |\hat{f}_{s,p} - 0.5|$ is large (at least 0.02 by default). Let $\mathcal{A}$ refer to the set of such samples. Observe that samples without substantial allelic imbalance are not informative for identifying haplotype switches.

2 Identify haplotype switches as the set $\mathcal{L}$ of bins $s$ where $\text{sign}(\hat{f}_{s-1,p} - 0.5) \neq \text{sign}(\hat{f}_{s,p} - 0.5)$ for *all* samples $s \in \mathcal{A}$.

3 Compute the proportion $|\mathcal{L}|/(N-1)$ of haplotype switches on the chromosome arm. If this proportion does not exceed a threshold (default 0.01), then do not correct haplotype switches.

4 If sufficiently many haplotype switches are detected, perform piecewise constant regression on the average haplotype frequencies across imbalanced samples. By default, we fit 10 segments to each chromosome arm.

5 Identify those segments that have a) sufficiently many haplotype switches (default $\geq 10\%$); b) sufficiently high average mhBAF (default $\geq 0.45$); and c) sufficiently high allelic imbalance in at least 1 sample (default $\geq 0.02$). For each such segment, assign haplotypes to each bin to minimize the pairwise difference in mhBAF between adjacent bins in the sample with the most extreme allelic imbalance (on average for the segment).

**Author details**

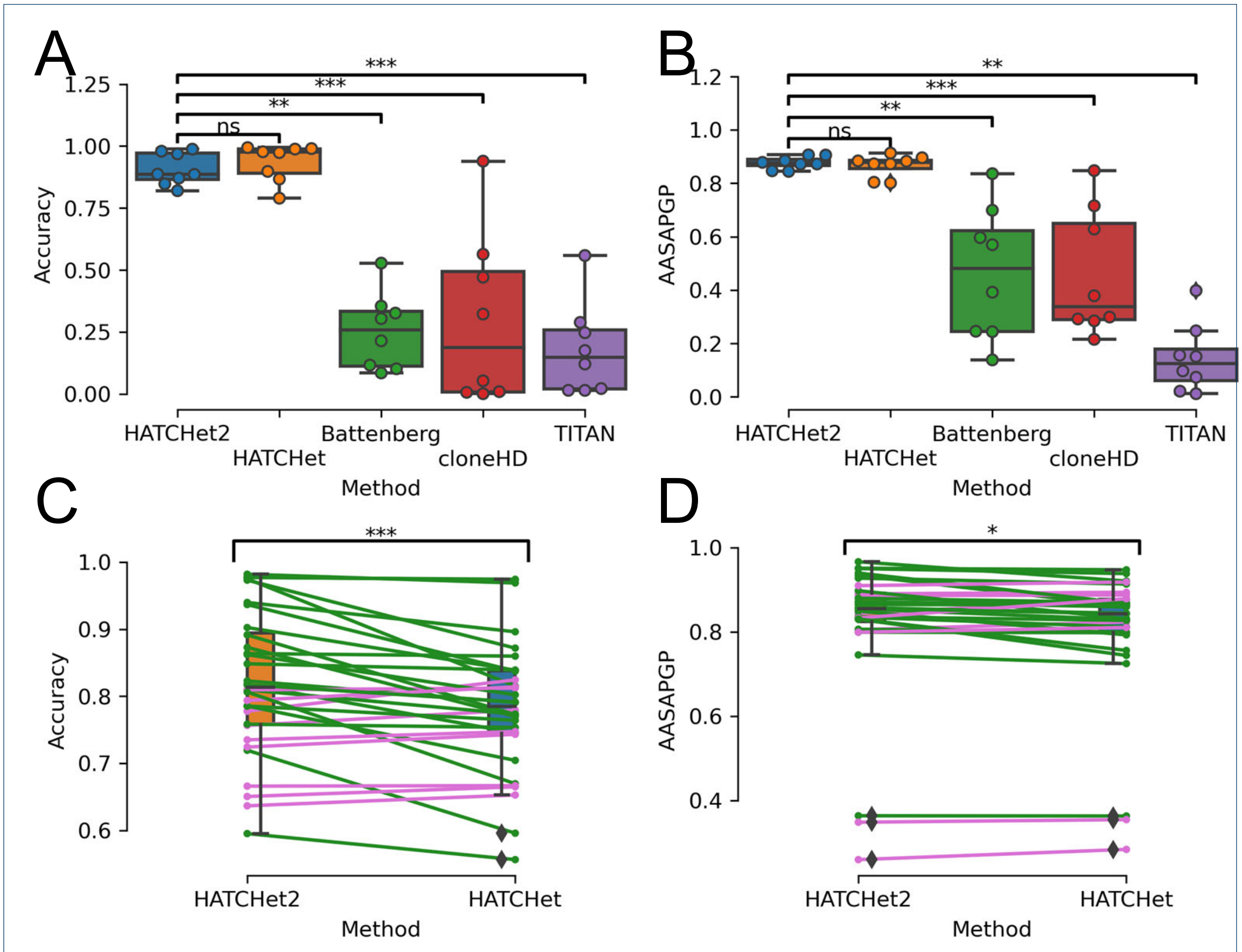

Figure S1: **Benchmarking HATCHet2, HATCHet [1], TITAN [4], cloneHD [3], and Battenberg [2] on simulated data**. **A-B** Results from all methods on 8 multi-sample datasets simulated using MASCoTE [1] measured by accuracy (A) and AASAPGP (B). For all panels, asterisks indicate significance by paired t-test: * $= 0.01 \leq p < 0.05$, ** $= 10^{-3} \leq p < 0.01$, *** $= p < 10^{-3}$. **C-D** Results from HATCHet2 and HATCHet [1] on the same data with each sample considered as an independent dataset ($n = 32$ samples), again measured by accuracy (C) and AASAPGP (D). Green lines indicate datasets on which HATCHet2 achieved a higher accuracy than HATCHet, whereas pink lines indicate that HATCHet achieved a higher accuracy.

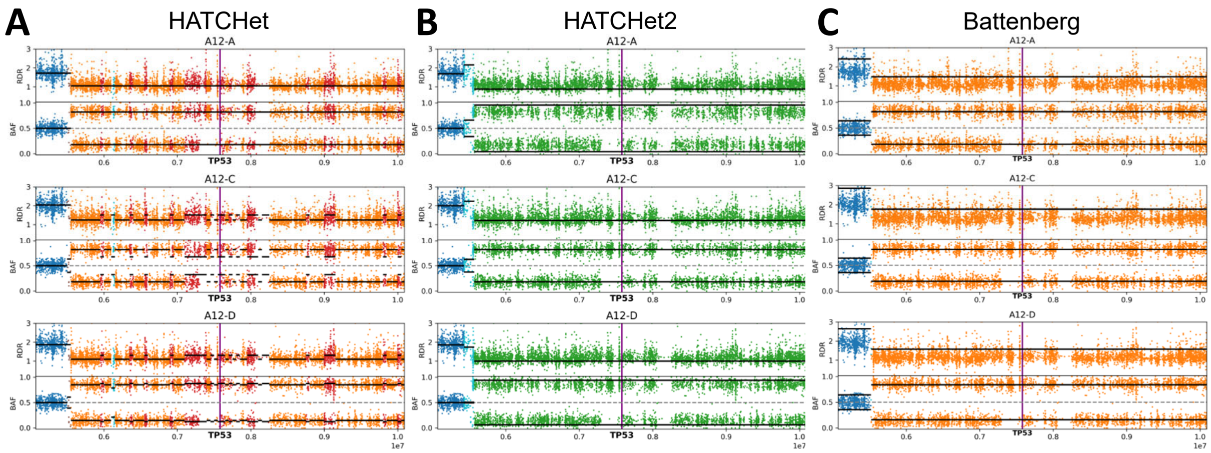

Figure S2: **Segments from HATCHet2, HATCHet [1], and Battenberg [2] near TP53 locus on chromosome 17 in patient A12**. As in Fig. 2, each point is a small region between adjacent SNPs (RDR) or a SNP (BAF) colored by the assigned copy-number state of the overlapping segment in the corresponding solution. Black bars indicate the expected RDR and BAF of each segment according to the copy-number states and mixture proportions identified by the corresponding method. Genes are represented as purple bars. **A** Solution from HATCHet. **B** Solution from HATCHet2. **C** Solution from Battenberg.

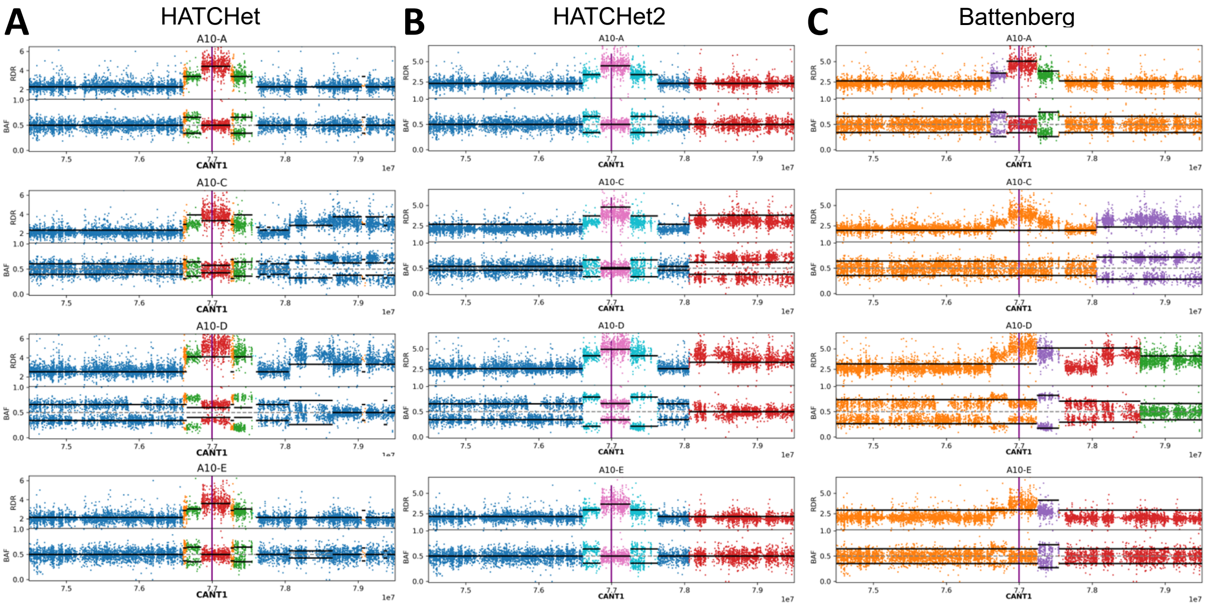

Figure S3: **Segments from HATCHet2, HATCHet [1], and Battenberg [2] near CANT1 locus on chromosome 17 in patient A10**. As in Fig. 2, each point is a small region between adjacent SNPs (RDR) or a SNP (BAF) colored by the assigned copy-number state of the overlapping segment in the corresponding solution. Black bars indicate the expected RDR and BAF of each segment according to the copy-number states and mixture proportions identified by the corresponding method. Genes are represented as purple bars. **A** Solution from HATCHet. **B** Solution from HATCHet2. **C** Solution from Battenberg.

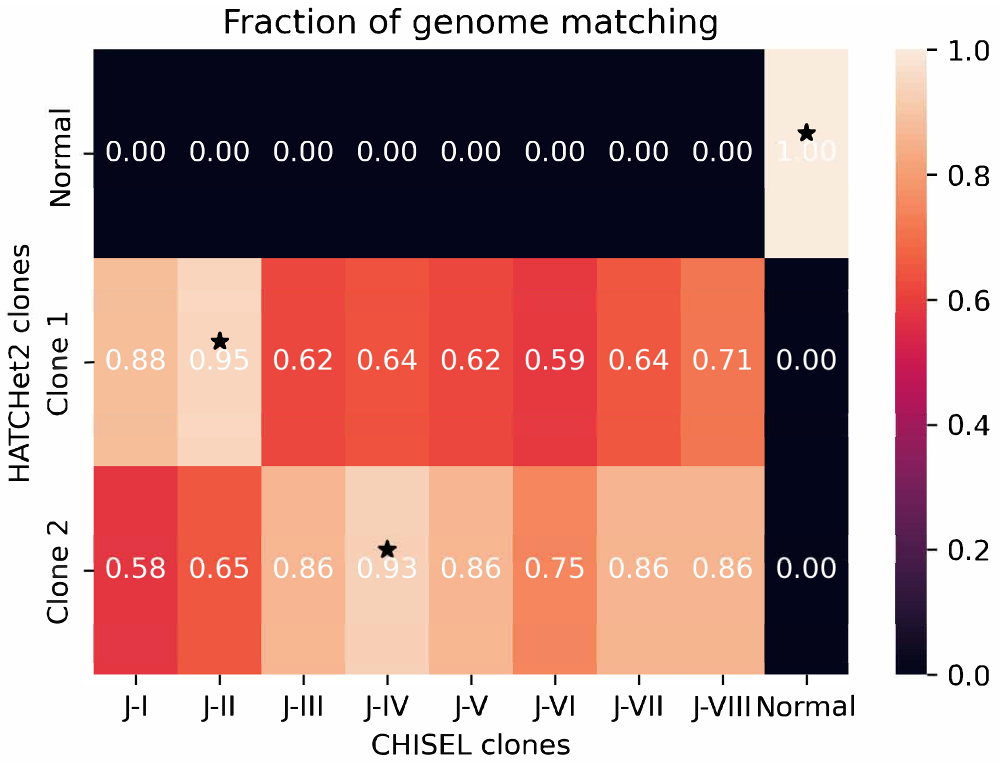

Figure S4: **Comparison between HATCHet2 clones and CHISEL [9] clones from 10X single-cell whole-genome sequencing data**. Proportion of the genome with the same haplotype-specific copy-number assignment between HATCHet2 clones (rows) inferred from pseudo-bulk samples and CHISEL clones (columns) inferred by aggregating single cells. Stars indicate the CHISEL clone which most closely matches each HATCHet2 clone.

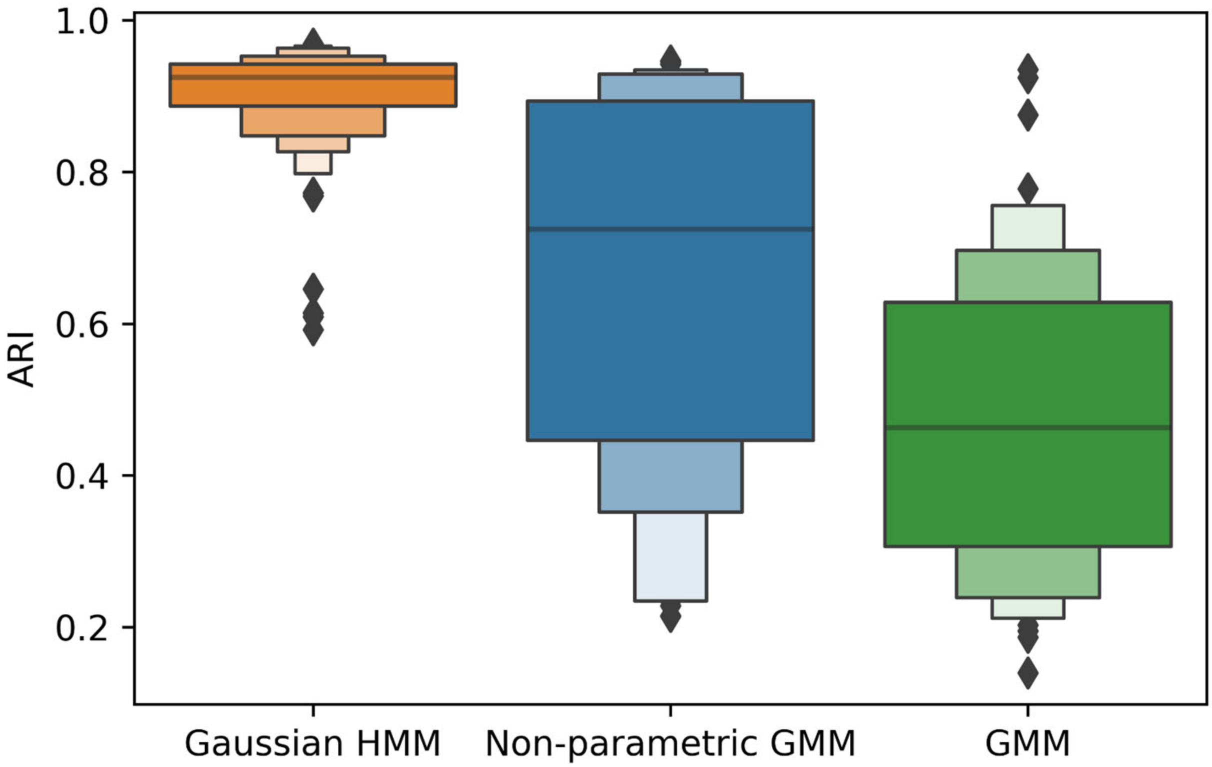

Figure S5: **Comparison of clustering methods on simulated data.** Comparison of clustering performance for 3 different methods in recovering distinct copy-number states from 64 simulated generated using MASCoTE [1]. Y-axis shows the adjusted Rand index (ARI) when comparing the inferred clusters to the ground truth copy-number events.

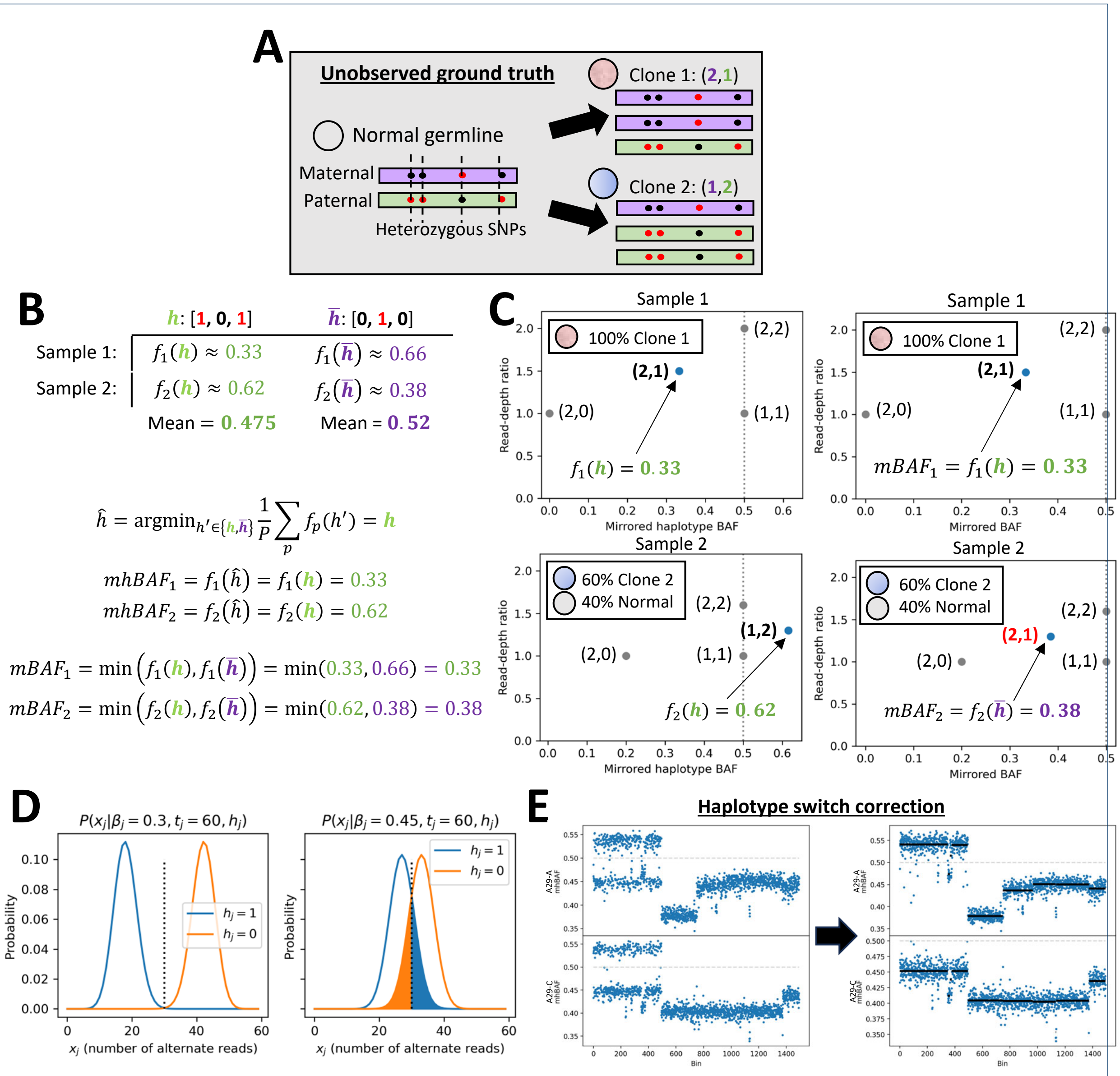

Figure S6: **Inference of minor haplotype BAF (mhBAF). A** Single-bin example of ground truth haplotypes and clone haplotype-specific copy-number profiles that we would like to infer. **B-C** Numerical example of the difference between the mirrored BAF (mBAF) and HATCHet2's mhBAF. The mBAF chooses between haplotypes in a sample-specific manner, unlike the mhBAF which measures the same haplotype across all samples. Panel B shows the computation of each statistic; panel C visualizes the two statistics in the context of theoretical values of haplotype-specific copy-number states. **D** Distributions of alternate counts $x_j$ for a single SNP $j$ (in a single sample) with different phasing $h_j$ and BAF $\beta_j$, under a simple binomial observation model. Shaded areas indicate regions where taking the $\arg\min$ would incorrectly assign the phase $h_j$ for SNP $j$. **E** Example of haplotype switch correction performed on chromosome 7 in two WGS samples from prostate cancer patient A29 from Gundem et al. [10]. Black bars indicate segments of similar average mhBAF identified by HATCHet2's correction algorithm.
